# Damage-induced senescent immune cells regulate regeneration of the zebrafish retina

**DOI:** 10.1101/2023.01.16.524296

**Authors:** Gregory Konar, Zachary Flickinger, Shivani Sharma, Kyle Vallone, Charles Lyon, Claire Doshier, William Lyon, James G. Patton

## Abstract

Zebrafish spontaneously regenerate their retina in response to damage through the action of Müller glia. Even though Müller glia (MG) are conserved in higher vertebrates, the capacity to regenerate retinal damage is lost. Recent work has focused on the regulation of inflammation during tissue regeneration with precise temporal roles for macrophages and microglia. Senescent cells that have withdrawn from the cell cycle have mostly been implicated in aging, but are still metabolically active, releasing pro-inflammatory signaling molecules as part of the Senescence Associated Secretory Phenotype (SASP). Here, we discover that in response to retinal damage, a subset of cells expressing markers of microglia/macrophages also express markers of senescence. These cells display a temporal pattern of appearance and clearance during retina regeneration. Premature removal of senescent cells by senolytic treatment led to a decrease in proliferation and incomplete repair of the ganglion cell layer after NMDA damage. Our results demonstrate a role for modulation of senescent cell responses to balance inflammation, regeneration, plasticity, and repair as opposed to fibrosis and scarring.

## Introduction

Degenerative eye diseases such as age-related macular degeneration (AMD), retinitis pigmentosa (RP), or Stargardt Disease affect millions of people each year. Each of these conditions is marked by profound atrophy and loss of one or more layers of the retina, which are unable to regenerate or recapitulate lost retinal neurons after damage. In contrast to humans and other mammals, zebrafish (*Danio rerio*) have the endogenous ability to regenerate their retina following damage ^1,2^. Damage to the zebrafish retina triggers a conserved mechanism of cellular regeneration driven by Müller Glia (MG), which serve as resident stem cells ^3–7^. After injury, MG dedifferentiate and undergo asymmetric division for both self-replacement and the generation of progenitor cells that proliferate to replace any lost neurons ^3^. Despite conservation of MG, regeneration is blocked in higher vertebrates. However, recent work and comparative analyses of zebrafish, chick, and mouse retina regeneration suggests that activation or de-repression of gene regulatory networks can enable limited reprogramming of mammalian MG ^8–12^. Mechanistic understanding of how zebrafish regenerate is crucial to discover novel factors that promote complete regeneration of the mammalian retina.

Recent papers have demonstrated a role for the immune system and inflammation in retina and retinal pigment epithelium regeneration ^13–19^. These effects appear to be mediated by resident and infiltrating macrophages and microglia that show dynamic transcriptional and morphological changes during regeneration ^20–24^. There are diverse populations of retinal microglia/macrophages, but the precise identity of distinct subsets of these cells throughout regeneration remains to be determined and it is not clear how changing patterns of secreted signaling molecules from these diverse subpopulations regulate the initiation and resolution of retina regeneration ^10,24–28^.

Senescence is a cellular process triggered by a myriad of molecular factors including aging, oxidative stress, telomeric shortening, DNA damage, inflammation, or epigenetic dysfunction ^29^. During senescence, cells exit the cell cycle in a programmed growth arrest event regulated by p53, p16 (Cdkn2a), and p21 (Cdkn1a) ^30–32^. Despite withdrawing from the cell cycle, senescent cells remain metabolically active and adopt a Senescence Associated Secretory Phenotype (SASP), which promotes the secretion of cytokines and growth factors that directly modulate the local cellular microenvironment ^33^,^34^. While senescence has been more well studied related to aging and cancer, it also plays a seemingly paradoxical role in development and regeneration ^35^. Transient initial exposure to pro-inflammatory SASP signaling appears to be required for regeneration, whereas chronic SASP exposure restricts the regeneration and proliferation of somatic or stem-like cells ^30^,^36^. Transient versus chronic senescent cell detection is apparent in models of spinal cord injury when comparing mammals and fish, where mammals show a chronic accumulation of senescent cells while fish show transient senescent cell expression ^37^. Targeting senescent cells with senolytic drugs (senotherapy) has been shown to aid in regeneration in multiple mammalian tissues including liver, kidney, muscle, and spinal cord ^38–42^.

In the retina, senescent cells are often associated with age related diseases such as AMD, and their targeting in retinal pigmented epithelial (RPE) cells has shown success in reducing AMD-like phenotypes in animal models of AMD ^43–46^. However, outside of the RPE, senescence of the inner retinal layers has seldom been investigated for its utility in contributing to regeneration of retinal neurons. Here, we provide the first evidence that in the adult zebrafish retina, there is a temporally conserved, transient senescent response to damage mediated by a subset of macrophages/microglia. Premature clearance of these senescent cells inhibits proliferation and leads to incomplete repair of the retinal ganglion cell layer after NMDA damage.

## Materials and Methods

### Zebrafish Husbandry and Maintenance

Wild-type AB zebrafish were maintained at 28.0°C on a 12:12hr light-dark cycle. All experiments were performed in accordance with Vanderbilt University Institutional Animal Care and Use Committee (IACUC) approval #M1800200.

### NMDA Damage Model

Adult zebrafish aged 5-10 months were anesthetized using a 0.016% Tricaine solution (MESAB, Acros Organics). The sclera of the left eye was cut using a sapphire blade, and 0.5uL of a 100mM NMDA solution was intravitreally injected. An equal volume of 1X PBS was used as an injection control for the NMDA damage.

### Pharmacologic treatments

For Metformin, a stock solution of 50mM Metformin (PHR1084, Millipore Sigma, US) was prepared in distilled water. Prior to use, the Metformin was diluted 1:500 in fish water to achieve a final working concentration of 100uM in the tank. Metformin water was changed daily to ensure consistent drug levels throughout the experiment. For ABT-263 (Navitoclax), a 30uM working solution of ABT-263 was made using 5% Tween-80, 20% PEG-300, and 2% DMSO in PBS, and 0.5uL were injected intravitreally.

### Immunohistochemistry, EdU labeling, and TUNEL assays

Fluorescent staining was performed as previously described ^47^. Briefly, zebrafish eyes were collected and fixed in 4% paraformaldehyde overnight at 4°C. Eyes were then moved to a sucrose gradient and stored overnight in 30% sucrose at 4°C. Following sucrose treatment, eyes were moved to 2:1 Cryomatrix:30% sucrose for 2 hours and embedded in Cryomatrix (Fisher Scientific). Sections were collected on a Leica cryostat using charged Histobond slides (VWR) at a thickness of 15um, then dried on a heating block and stored at −80°C. For IHC, slides were warmed and then incubated in 1X PBS for 30 minutes to rehydrate. Slides were then subjected to antigen retrieval as previously described ^48^, before cooling to room temperature. Blocking solution (3% donkey serum, 0.1% Triton X-100 in 1X PBS) was applied to the slides and incubated for 2h at room temperature (RT) before incubation with primary antibodies overnight at 4°C. The antibodies used for IHC are listed in **Table 1**.

| <u>Name</u> | <u>Target</u> | <u>Concentration</u> | <u>Source</u> |
| --- | --- | --- | --- |
| Proliferating Cell Nuclear Antigen (PCNA) | Actively proliferating cells | 1:500 | Abcam |
| HuC/D | Retinal Ganglion Cells | 1:500 | Invitrogen |
| 4c4 | Macrophages/Microglia | 1:1000 | Hitchcock Lab (Univ. Michigan) |
| Lcp1 | Macrophages/Microglia | 1:500 | GeneTex |

After primary antibody incubation, slides were washed and stained with secondary antibodies in 1% Donkey Serum, 0.05% Tween-20 in PBS along with 1:1000 To-Pro-3 (Invitrogen) for 2h at RT. Secondary antibodies used include donkey anti-mouse 488 (1:500), donkey anti-rabbit 488 (1:500), donkey anti-mouse cy3 (1:500), and donkey anti-rabbit cy3 (1:500) (all Jackson Immuno). Slides were then washed, mounted, and coverslipped using VectaShield Antifade Mounting Medium (Vector Labs).

For EdU labeling, adult zebrafish received a 0.5uL intravitreal injections of 10mM EdU at both 24h post damage (hpi) and 48hpi. Incorporation was detected using the Click-iT^™^ Plus EdU Cell Proliferation Kit (555 fluorophore, ThermoFisher, US). For TUNEL staining, a Click-iT^™^ Plus TUNEL Assay Kit was used (488 fluorophore, ThermoFisher, US).

### Senescence associated b-galactosidase staining

Senescence associated b-galactosidase (SA-βGal) staining was performed on retina sections using a kit from Cell Signaling Technology. Briefly, slides were warmed on a heating block and incubated in PBS. SA-βGal staining solution was warmed to 37°C with the pH adjusted to between 5.9-6.1 and then applied to retinal sections and incubated overnight at 37°C. Samples were then washed and incubated with a 1:1000 dilution of To-Pro-3 (ThermoFisher) in PBS for nuclear staining and then mounted using VectaShield antifade mounting medium (Vector Labs) and sealed using clear nail polish.

### Imaging and Image Processing

For imaging of IHC, EdU, or TUNEL staining, a META Zeiss LSM Meta 510 confocal microscope was used through the Vanderbilt Center for Imaging Shared Resource (CISR) Core. Slides stained with SA-βGal and ToPro were imaged using a Nikon AZ100M Widefield microscope through the Vanderbilt CISR Core. Confocal images were processed using Zen Blue version 3.1 with follow up analyses done using ImageJ. AZ100M images were first processed using NIS-Elements Viewer 5.21 and then subsequently analyzed using ImageJ.

### Quantification and Statistical Analysis

Experiments involving immunostaining and cell number quantification were evaluated in an unbiased and blinded manner. Briefly, stained slides that comprised the optic nerve region were evaluated in the central retina. Both dorsal and ventral regions were counted to encompass the entire retinal region, spanning approximately ~300-400mm from the optic nerve in either direction. Regions of extreme structural damage or nearer to the ciliary marginal zone (CMZ) were excluded from counts as to not cause artificially elevated numbers. Fluorescent cell counts from two independent sections were counted and averaged from each eye. For gap counting, slides were stained with HuC/D primary antibodies and To-Pro-3 nuclei marker and regions of missing retinal ganglion cells were counted under a fluorescent microscope. Significance was calculated using a two-way ANOVA with Tukey’s post-hoc test for inter-group comparisons in GraphPad Prism 9.4.1. All data represented in graphical form is of the average ± standard error of the mean (s.e.m). The number of fish used in each experiment is described in the figure legends.

## Results

### Detection of senescent cells after NMDA damage

Traditionally, senescence has been viewed as largely associated with aging, but recent evidence supports a role for senescence in development and regeneration ^35,49,50^. After spinal cord damage, senescent cells can be transiently detected in zebrafish whereas in mice, senescent cells accumulate over time and targeting of senescent cells in mice with senolytic drugs can improve tissue repair ^37^. We sought to determine if senescent cells are detectable in the regenerating retina, whether they follow a similar burst and clearance profile as in the zebrafish spinal cord damage model, and whether temporal treatment with senolytics might affect repair. To first determine whether we could detect senescent cells, we damaged zebrafish retinas by intravitreal injection of N-methyl-D-Aspartate (NMDA) which causes excitotoxic damage of retinal ganglion cells (**Figure 1A**) ^8^. We then stained tissue sections for Senescence Associated b-Galactosidase (SA-βGal), a lysosomal marker of senescent cells ^51–53^. Staining of both damaged and undamaged retina sections across an 18-day time course detected the presence of senescent cells expressing SA-βGal in the damaged retina (**Figure 1B**). Senescent cells were first detectable in NMDA damaged retinas at 2 days post injury (dpi) (p<0.05) and then began to decline after 12 dpi (p<0.05). Senescent cells were largely undetectable in undamaged or PBS injected eyes. Detection of senescent cells was not limited to the ganglion cell layer (RGC), as senescent cells were observed in all retinal layers spanning from the outer nuclear layer (ONL) to the RGC layer (**Figure 1C**). The expression pattern and timing of appearance of senescent cells in the inner retina is consistent with the temporal appearance and clearance of SA-βGal+ cells after spinal cord damage in zebrafish and stands in contrast to senescent cell accumulation in the mammalian retina or RPE after damage ^37,43,54,55^. Reduced detection or clearance of senescent cells as regeneration proceeds may constitute a key difference between regenerative responses in zebrafish compared to higher vertebrates.

**Figure 1.**
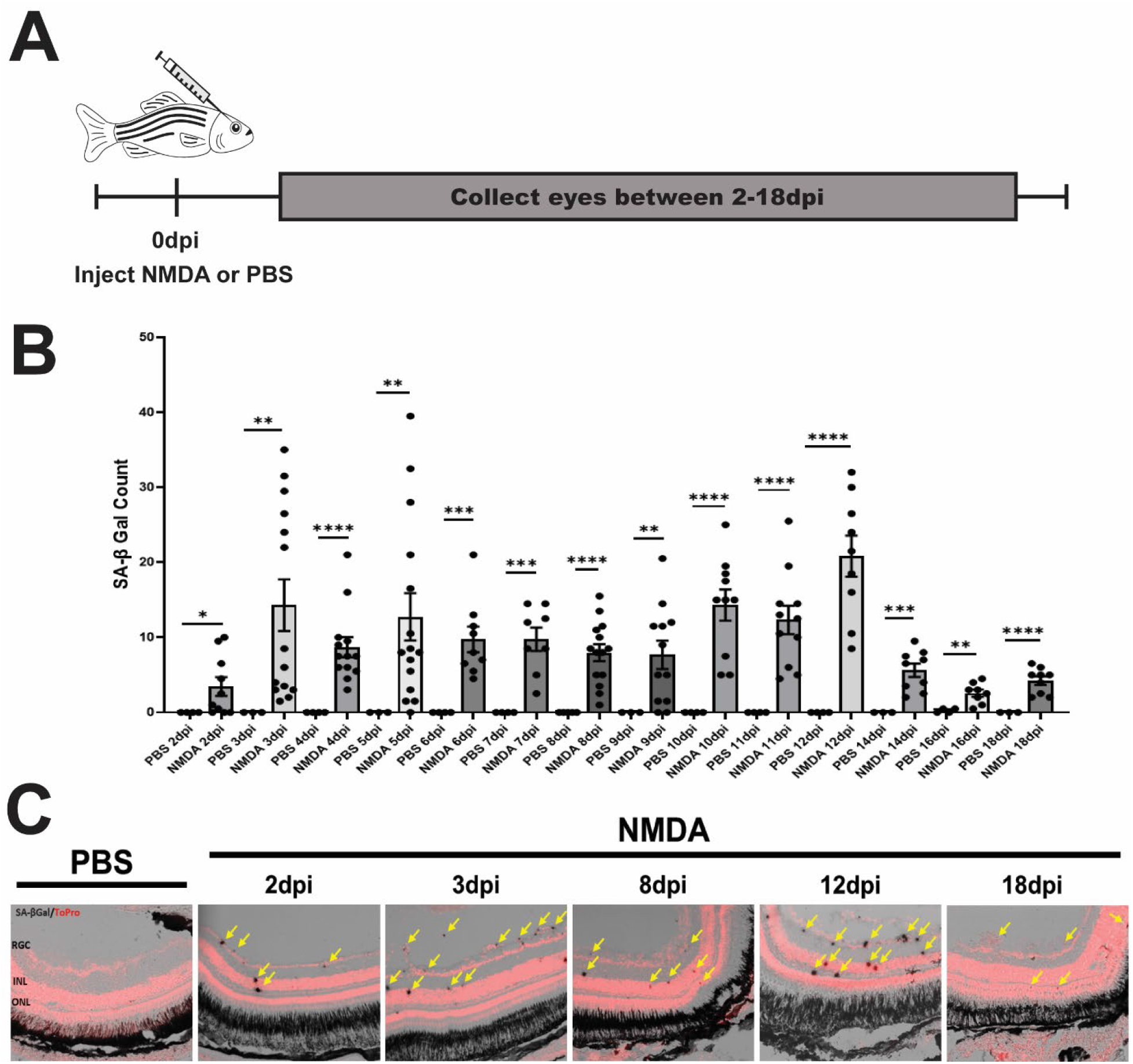
Detection of senescent cells in the zebrafish retina after NMDA damage. (A) Schematic highlighting experimental timeline. Wildtype AB adult zebrafish aged 5-10 months were intravitreally injected with either 0.5uL of 100mM NMDA or PBS and then eyes were collected between 2-18 days post injection (dpi) for analysis. (B) Eyes injected with NMDA or PBS were sectioned and stained for Senescence Associated b-galactosidase (SA-βgal) expression. Senescent cells were detected beginning at 2dpi (n=10) followed by declining numbers after 12dpi (n=9). Undamaged or PBS control eyes were largely devoid of senescent cells except for a few puncta at 16dpi (n=4). (C) Representative images of SA-βgal stained retina sections. Nuclear staining was performed using To-Pro-3 in red, SA-βgal cells are in dark grey. ONL=Outer Nuclear Layer. INL=Inner Nuclear Layer. RGC=Retinal Ganglion Cell Layer. All graphs show individual eyes with mean ± s.e.m. All statistics done using Student’s T-test where * = p<0.05, ** = p<0.01, *** = p<0.001, and **** = p<0.0001.

### Macrophages/microglia become senescent after retina damage

We next sought to determine the origin and identify of the SA-βGal+ cells that we detected after NMDA damage. For this, we immunostained sections after NMDA damage with antibodies against L-plastin (Lcp1) which marks macrophages and microglia ^20,21,56^. We discovered that a distinct subset of macrophages/microglia that are marked by expression of Lcp1 co-localize with SA-βGal+ cells (**Figure 2A**). The Stenkamp and Mitchell labs have shown that after retina damage in the zebrafish induced by injection of the neurotoxin oubain, immune cells, including resident microglia and infiltrating macrophages, rapidly respond with morphological changes and altered gene expression ^20,21^. Further, depletion of microglia and macrophages inhibits retina regeneration in both the chick and in zebrafish ^24,57^. Quantitation of the relative levels of Lcp1+ and SA-βGal+ cells showed that approximately 30% of Lcp1+ cells co-localize with SA-βGal+ (**Figure 2B**) demonstrating that after NMDA damage, the overall immune response involves both macrophage/microglial responses and a senescent cell response.

**Figure 2.**
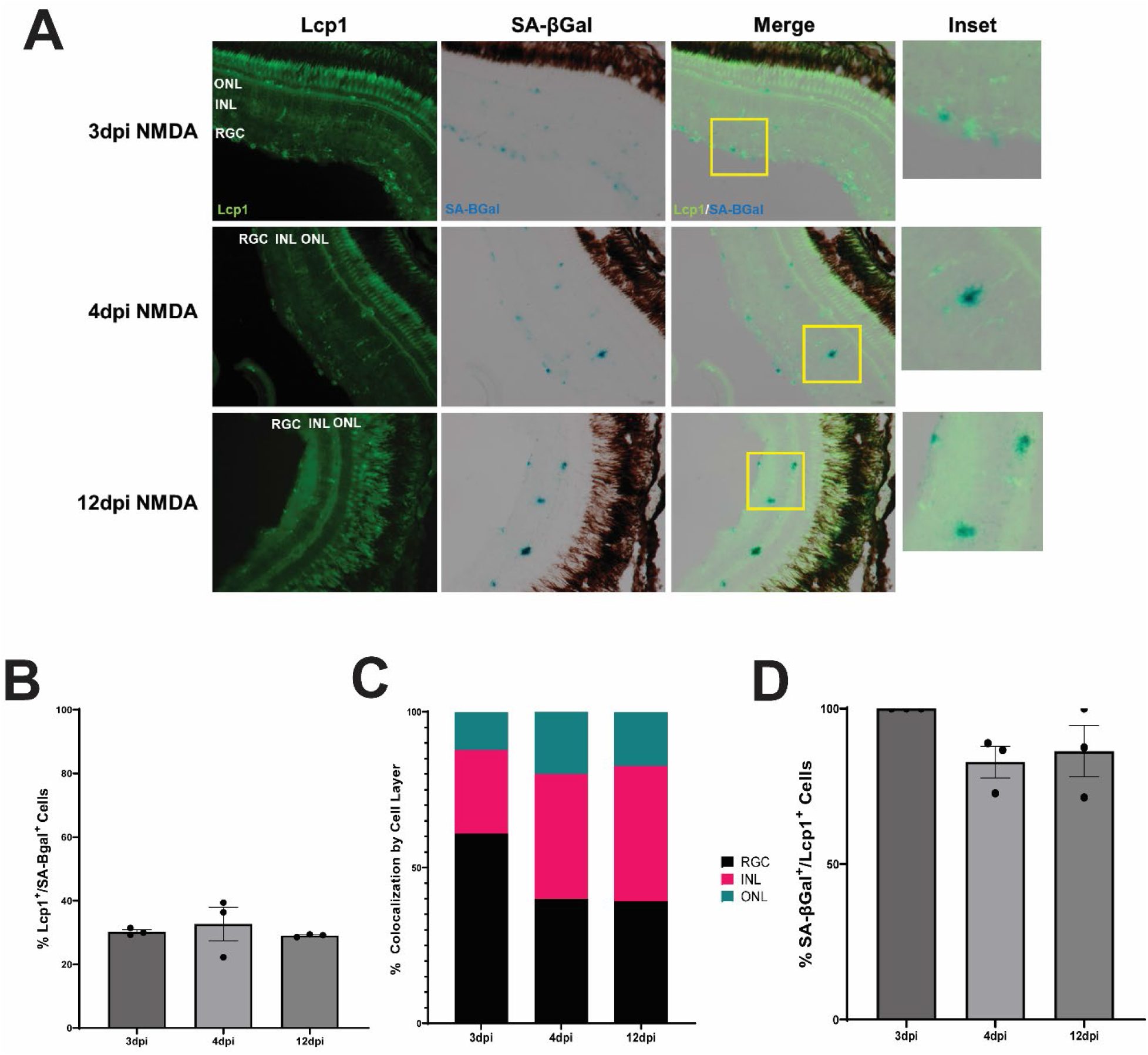
Co-localization of L-plastin and SA-βgal after NMDA damage. (A) NMDA treated retina sections from 3, 4, or 12dpi were immunostained for L-plastin (Lcp1), a marker of macrophages/microglia, and for SA-βgal. Damaged retinas showed co-localization of Lcp1 (green) and SA-βgal (teal) at all timepoints (see inset). (B) Graph showing the percentage of Lcp+ microglia/macrophages that were also SA-βgal+ in the retina across 3, 4, and 12dpi. Only a small subset of Lcp1+ cells showed SA-βgal+ staining, and the percentage was not statistically different across the 3 timepoints measured (p>0.05). (C) Overall percent distribution of Lcp1+/ SA-βgal+ cells in the indicated retinal layer at each timepoint evaluated. (D) Graph showing the percentage of SA-βgal+ cells that were also Lcp1+ at each timepoint. The majority of SA-βgal+ co-localized with Lcp1+ cells across all timepoints, and the percentage was not statistically different at any point (p>0.05). All data are the product of n=3 retinas. ONL=Outer Nuclear Layer. INL=Inner Nuclear Layer. RGC=Retinal Ganglion Cell Layer.

The majority of Lcp1^+^/SA-βGal^+^ cells were detected in the RGC layer, but dual labeled cells were observed across all three layers of the retina (**Figure 2C**). To assess whether all SA-βGal^+^ cells co-localize with microglia/macrophages or whether other cell types become senescent after damage, we quantified the total number of SA-βGal+ cells in each retinal section and divided by the total number of Lcp1^+^/SA-βGal^+^ cells. We found that at 3dpi, 100% of the SA-βGal^+^ cells were also Lcp1^+^/SA-βGal^+^ cells, and that at 4dpi and 12dpi, 83% and 87% SA-βGal^+^ cells overlap with Lcp1^+^ cells (**Figure 2D**). These data suggest that the primary senescent cell type that is detectable in response to damage is derived from microglia/macrophages ^15,16^.

### Senotherapeutic drugs reduce proliferation after NMDA damage and inhibit regeneration

The link between senescence and aging has prompted the development of drugs to clear senescent cells. We tested the effects of two different senotherapeutic drugs on retina regeneration, ABT-263 and Metformin. ABT-263 (Navitoclax) is a Bax-Bcl2 inhibitor that clears senescent cells through stimulation of apoptosis ^58,59^. Metformin is a common anti-diabetic drug that targets multiple aging and cancer pathways and has been shown to function as a senolytic agent through the stimulation of autophagy ^60–63^. Interestingly, senotherapeutics, especially Metformin, are being used to treat age-related macular degeneration ^54^.

Treatment with Metformin in NMDA treated retinas (**Figure 3A**) led to a marked decrease in SA-βGal^+^ cells in the retina at 5 and 10 dpi, confirming that Metformin functions as a senolytic (**Figure 3B,C**). ABT-263 treatment also reduced the number of senescent cells in NMDA treated retinas between 5-10dpi (**Figure 3 D,E**). Finally, combination treatment of both Metformin and ABT-263 also reduced the number of senescent cells in NMDA treated retinas between 5-10dpi (**Figure 3 F,G**). These results confirm that senolytic treatment leads to a decrease in senescent cell numbers and that damage-induced appearance of senescent cells after retina damage can be modulated by senolytic treatment.

**Figure 3.**
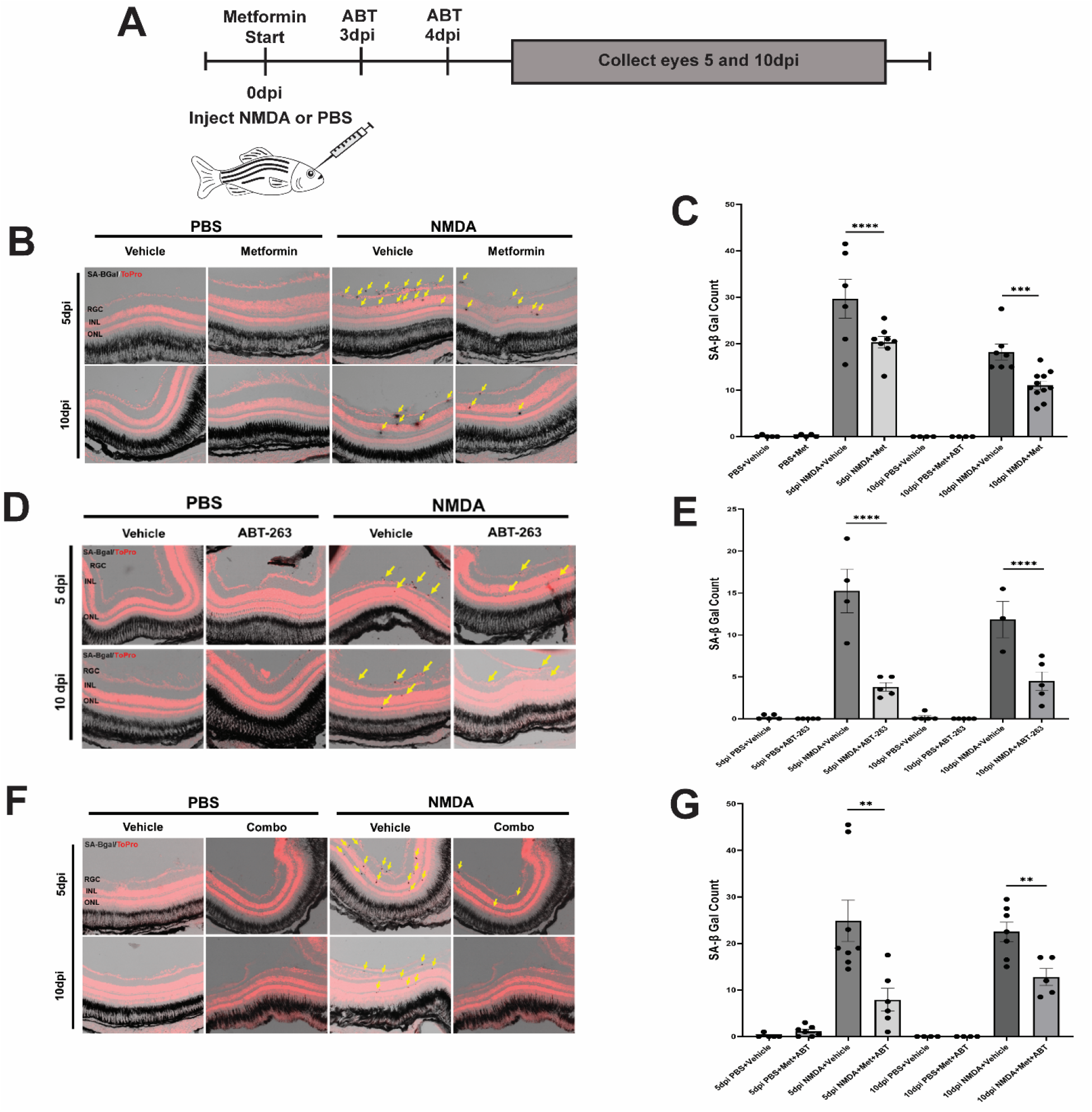
Senolytic treatment decreases SA-βgal+ cells in NMDA damaged retinas. (A) Schematic depicting experimental timeline of senolytic dosing in conjunction with NMDA damage. Adult wildtype (AB) fish were intravitreally injected with 0.5uL of 100mM NMDA or PBS, and then placed in tanks containing either 100uM Metformin or standard tank water. Fish receiving ABT-263 injections received 0.5uL intravitreal injections of 30uM ABT-263 on days 3 and 4 after initial NMDA injection. (B-C) Metformin treatment reduced the number of senescent cells in NMDA treated retinas between 5-10dpi (n=5-11). (D-E) ABT-263 treatment reduced the number of senescent cells in NMDA treated retinas between 5-10dpi (n=3-5). (F-G) Combination treatment of Metformin and ABT-263 additionally reduced the number of senescent cells in NMDA treated retinas between 5-10dpi (n=4-8). ONL=Outer Nuclear Layer. INL=Inner Nuclear Layer. RGC=Retinal Ganglion Cell Layer. All graphs show individual eyes and mean ± s.e.m. All statistics were performed using Student’s T-test where * = p<0.05, ** = p<0.01, *** = p<0.001, and **** = p<0.0001

### Senolytic treatment limits cellular proliferation in NMDA damaged retinas

Following retinal or brain injury in zebrafish, an initial inflammatory response is required to initiate regeneration ^13–15,19,24,64^. MG respond to inflammatory signals from dying neurons ^65^ and from resident and infiltrating macrophages and microglia ^14,19–21,24^. Senescent cells release cytokines and other inflammatory signals that could increase proliferation of immune cells (and MG) during the initial stages of retina regeneration. To test whether senolytic clearance of SA-βGal^+^ cells affects proliferation in the damaged retina, we injected EdU at various times after injury in the presence or absence of ABT-263, Metformin, and the combination thereof, followed by EdU staining of sections at 5 and 10dpi. All three senoytic treatments showed modest but signifcant decreases in proliferation as measured by EdU incorporation (Metformin, ABT p<0.05., Combo p<0.01) (**Figure 4**). These results support the hypothesis that senescent cells mediate signaling events that induce proliferation after retina damage.

**Figure 4.**
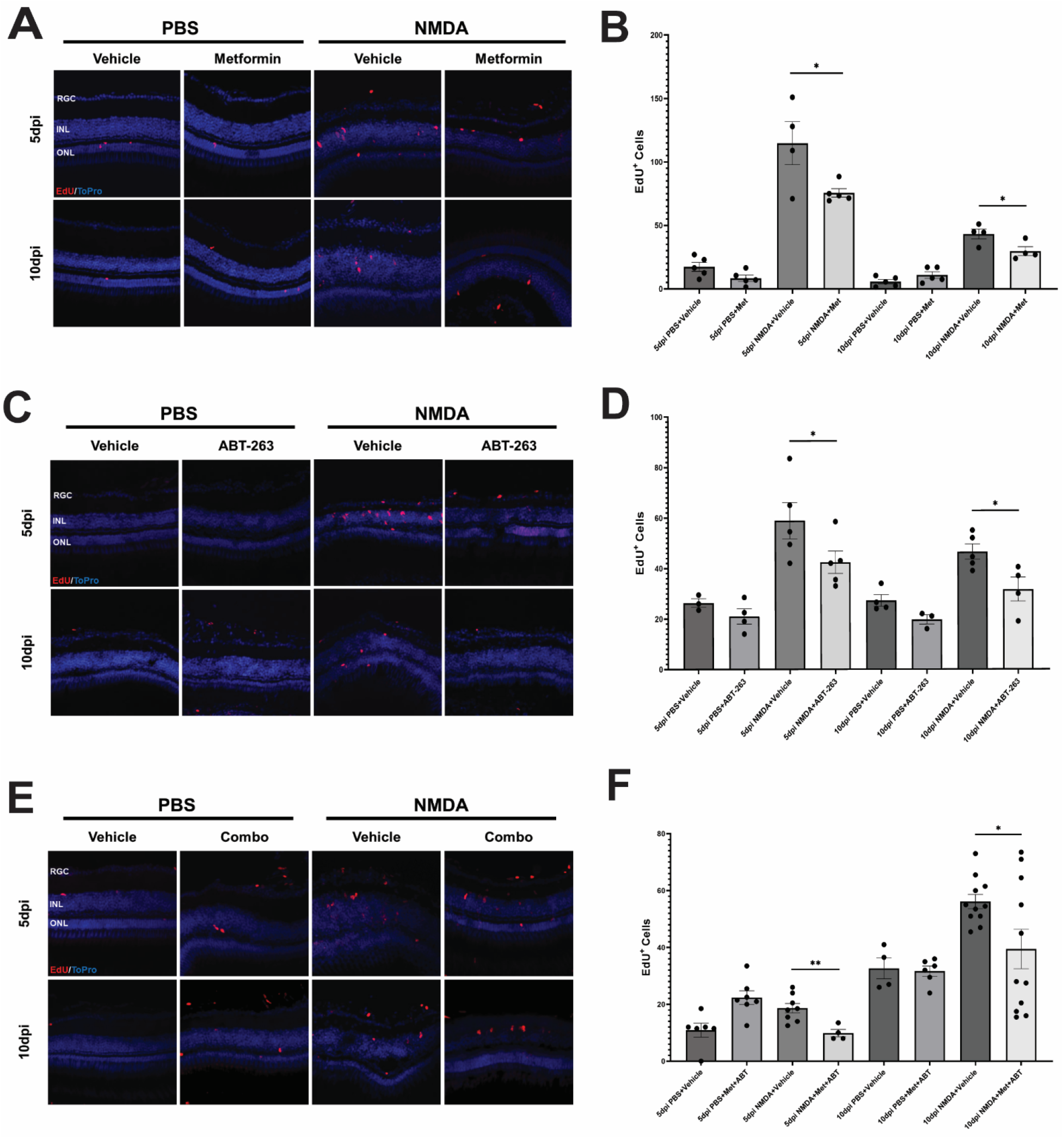
Senolytic treatment limits proliferation in NMDA damaged retinas. (A-B) Wildtype AB zebrafish received either 0.5uL intravitreal injections of 100mM NMDA or PBS, were placed in Metformin-treated or regular tank water, and were then intravitreally injected with EdU at the indicated time points. Metformin treatment showed a significant decrease in the number of EdU+ cells in NMDA treated retains (n=4-5). (C-D) Wildtype zebrafish were intravitreally injected with ABT-263 three and four days after NMDA injection. ABT-263 treatment produced a significant decrease in EdU+ cells in NMDA treated retinas at both 5 and 10dpi (n=3-5). (E-F) Combination treatment with Metformin and ABT-263 showed a decrease in EdU+ cells in NMDA treated retinas at 5 and 10dpi (n=4-12). ONL=Outer Nuclear Layer. INL=Inner Nuclear Layer. RGC=Retinal Ganglion Cell Layer. All graphs show individual eyes and mean ± s.e.m. All statistics were performed using Student’s T-test where * = p<0.05, ** = p<0.01, *** = p<0.001, and **** = p<0.0001

Since resident and infiltrating immune cells can be detected after retinal damage, we sought to determine if senolytic treatment impacted the overall number of 4c4^+^ immune cells in the retina. The 4c4 antibody targets the galectin 3 antigen in zebrafish microglia ^66^. All three senolytic treatments showed little to no change in 4c4^+^ cells in the retina (**Figure S1–3**). These results indicate that the decrease in proliferation observed in the retina after senolytic treatment is not due to significant depletion of microglia or macrophage based immune cells.

### Premature removal of senescent cells impairs RGC regeneration

We next sought to test whether senescent cells play a role in retina regeneration by determining whether clearance of senescent cells using senolytics would affect regeneration after NMDA damage. For this, we immunostained retinas treated with NMDA at 10dpi with antibodies against HuC/D, a marker of RGCs and amacrine cells ^67^. Without senolytic treatment, successful repair of NMDA damage in the RGC layer shows only a small number of gaps devoid of both nuclei and RGC staining at 10dpi (**Figure 5**). In contrast, the addition of ABT-263, Metformin, or the combination thereof, led to a significant increase in the number of gaps in the RGC layer (**Figure 5**). Compared to the control, the increase in gaps that alter the typical morphology of the RGC layer indicates incomplete regeneration of ganglion cells following NMDA damage and supports a role for senescent cell signaling to properly modulate regenerative responses.

**Figure 5.**
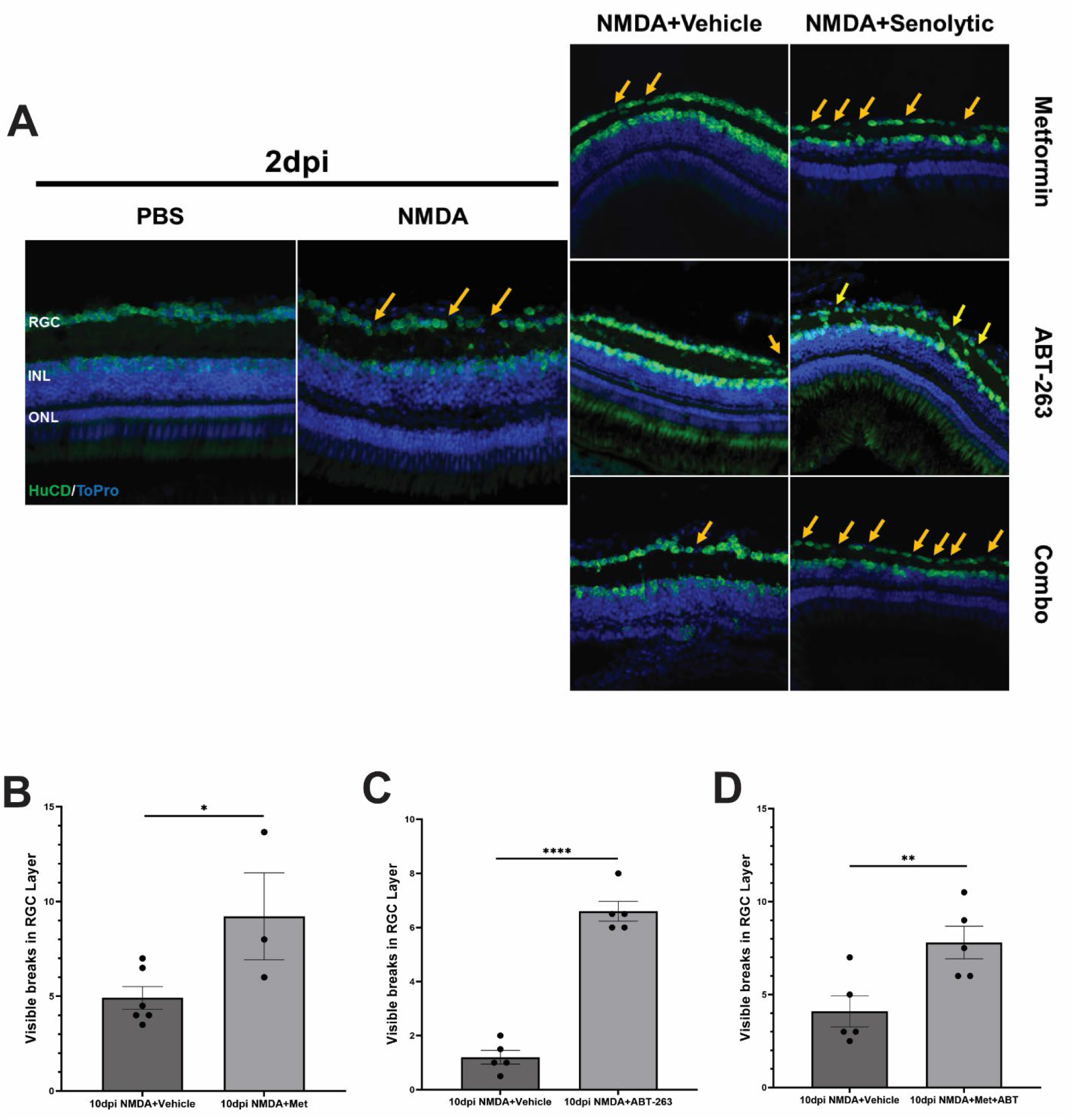
Premature removal of senescent cells impairs RGC regeneration in NMDA damaged retinas at 10dpi. (A) Wildtype AB zebrafish were intravitreally injected with 0.5uL of 100mM NMDA or PBS, and then subjected to either Metformin, ABT-263, or the combination. Representative images of PBS and NMDA treated retinas at 2 and 10dpi showing gaps in the RGC layer. (B-D) Quantification of gaps and breaks in the RGC layer. All data are the product of between 3-6 retinas. ONL=Outer Nuclear Layer. INL=Inner Nuclear Layer. RGC=Retinal Ganglion Cell Layer. All graphs show individual eyes and mean ± s.e.m. All statistics were performed using Student’s T-test where * = p<0.05, ** = p<0.01, *** = p<0.001, and **** = p<0.0001

We additionally sought to determine if the breaks in the RGC layer coincide with an increase in the number of apoptotic cells in damaged retinas. For this, we immunostained retinas treated with NMDA and the combination of both senolytic agents followed by TUNEL assays to identify apoptotic cells. With the addition of NMDA, we observed the expected increase in the number of TUNEL^+^ cells throughout the retina. However, we observed a further increase in the number of TUNEL^+^ cells after the combination of Metformin and ABT-263 treatment (p<0.05) (**Supp. Figure 4A-B**). These TUNEL^+^ cells were observed across all layers of the retina, consistent with the widespread localization of senescent cells after NMDA damage (**Figure 2**). Thus, in conjunction with decreased proliferation and impaired regeneration after senolytic treatment, there is a concomitant increase in cell death that is consistent with clearance of senescent cells.

## Discussion

Here, we show that following NMDA damage, senescent cells are detectable during zebrafish retina regeneration. Senescent cell detection after retina damage in zebrafish is transient, with a progressive accumulation of senescent cells beginning around 2-3 dpi and then decreasing as regeneration proceeds from 12-18 dpi. We also show that the retinal cells that display expression of SA-βGal co-localize with markers of macrophages and microglia. Lastly, we show that premature removal of immune-derived senescent cells via senolytic treatment leads to decreased proliferation following NMDA damage and incomplete regeneration of the retinal ganglion cells with increased breaks and gaps in the RGC layer.

Recent results support a modulatory role for inflammation during retina regeneration ^13–19,24^. We show for the first time that senescent cells play a role in the overall inflammatory response during retina regeneration, consistent with roles for senescent cells in plasticity and regeneration ^35,49,50^. Because we observed a decrease in proliferation after senolytic treatment, but not an overall decrease in the numbers of microglia/macrophages, it seems that signaling from senescent cells affects MG-derived proliferation which further supports a role for inflammatory signaling during retina regeneration. Beyond the retina, senescent cells have been shown to play a role in fin and spinal cord regeneration in zebrafish ^37,41^. Together, the data support a conserved and important role for senescent cells during retina regeneration. We refer to the senescent cells that are detectable after NMDA treatment of the retina as Damage Induced Senescent Immune (DISI) cells (**Figure 6**).

**Figure 6.**
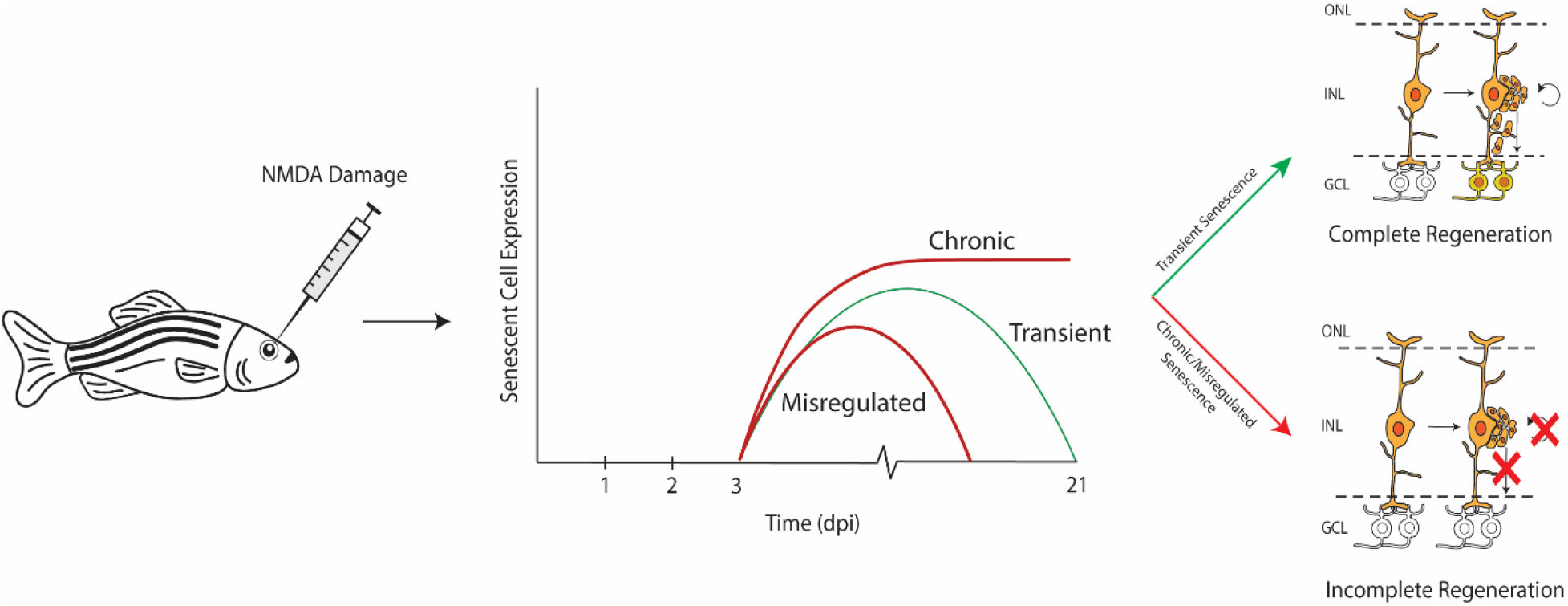
Senescence and zebrafish retina regeneration. After damage to the retina, senescent cells were detectable beginning at 2-3dpi and then decreasing after 12dpi. The transient presence of senescent cells is proposed to modulate paracrine and autocrine signaling to allow proper regeneration. Having too much (chronic) or too little (misregulated) signaling from senescent cells is proposed to result in an overall truncation of the regenerative response.

### Inflammation and Regeneration

Zebrafish possess the innate ability to regenerate any part of their body ^68^. Understanding the surrounding microenvironmental cues and cellular factors that facilitate this capability is important for complete understanding of regeneration and the role that such signaling may play in different mechanisms of regeneration, such as blastema formation in fin regeneration and MG-derived retina regeneration. During MG-derived retina regeneration the data support a model in which early pro-inflammatory responses give way to anti-inflammatory responses as regeneration proceeds ^13–17,69^. Our detection of immune cells expressing SA-βGal starting at approximately 2-3 dpi corresponds to similar timing for the transition from pro-to anti-inflammatory cytokine signatures during retina regeneration ^14^.

A senescent cell burst during the early stages of regeneration followed by a later decline in senescent cells is consistent with a role in modulating inflammatory response through the senescence associated secretory phenotype (SASP). Despite withdrawing from the cell cycle, senescent cells remain metabolically active ^70,71^. Secreted metabolites by senescent cells include cytokines, proteases, and extracellular matrix (ECM) components ^72^. One prominent cytokine associated with SASP signaling is interleukin 6 (IL-6), a cytokine that works to control localized inflammatory responses ^73–75^. Interestingly, IL-6 displays both pro- and anti-inflammatory roles ^76^ and its release by senescent cells is critical for cellular reprogramming ^77^. While it remains to be seen what additional factors are being secreted by senescent cells in the retina and any temporal changes in such factors, the shift from a proinflammatory environment championed by microglia and macrophages into a more anti-inflammatory or pro-regenerative environment correlates with the temporal appearance and clearance of senescent cells that we observe during retina regeneration. This is also consistent with results from spinal cord repair experiments in mice and zebrafish where senescent cells are cleared in regeneration competent zebrafish but persist and even continue to increase post spinal cord injury in mice ^37^.

Secondary to cytokine release from senescent cells is the release of various growth factors including granulocyte macrophage colony-stimulating factor (gmCSF) and matrix metalloprotease 9 (mmp-9) ^17,78^. GmCSF promotes retina regeneration ^78^ while mmp-9 is upregulated in both senescent cells and MG-derived progenitors and plays a role in modulating inflammation during early and late stages of photoreceptor regeneration ^17^. This highlights the ability of SASP factors to influence the microenvironment through paracrine and autocrine signaling that in turn modulates the responses of both immune cells and MG. Initial upregulation of anti-inflammatory factors followed by later secretion of pro-regenerative factors appears to provide the precise microenvironmental context for promoting regeneration after damage in the adult zebrafish retina.

### Senolytics and Regeneration

Senolytic treatment has the potential to serve as a therapeutic mechanism for many different diseases affecting a variety of organs including the eye ^43^, spinal cord ^37^, liver ^79^, as well as cancer, and other age-related diseases ^80–82^. Most experiments examining senescence and the use of senolytics have been performed in mammalian models where there is a distinct lack of senescent cell clearance over time ^37^. Removal of senescent cells via senolytic treatment has the potential to alleviate degenerative diseases ^83,84^. In the zebrafish retina, we show that senescent cell clearance occurs naturally and is temporally controlled. Zebrafish retina regeneration is generally completed within 3 weeks post injury ^1,7^, and the clearance of the senescent cells we observed is consistent with this time frame. Premature clearance of senescent cells caused incomplete repair indicating temporal mechanisms regulating the appearance and clearance of senescent cells, which we hypothesize coincides with the switch from pro-inflammatory to pro-regenerative responses.

### Inflammation and Regeneration in Mice and Zebrafish

Work in zebrafish retina regeneration has shown a role for initial inflammatory responses and dynamic changes in microglia and macrophages ^14–16,20,21,24^. In contrast, work in mice indicates that microglia suppress retina regeneration ^10^. This could indicate that mechanistic differences in inflammatory signaling underlie the ability to regenerate when comparing fish and mice. Alternatively, it is possible that temporal and context-specific differences can explain apparent contradictions between fish and mice regarding a role for immune cells and inflammation ^19^. From spinal cord regeneration experiments ^37^ and our work here, it seems that the ability to clear senescent cells is crucial for regeneration.

In summary, our results demonstrate that in response to NMDA induced ganglion cell damage, there is a population of senescent cells that display a temporal pattern of appearance and clearance as regeneration proceeds. These cells appear to be a subset of macrophages and microglia consistent with precise control of inflammation during regeneration. Finally, inducing premature clearance of these senescent cells through either autophagic or apoptotic mechanisms, showed a surprising decrease in proliferation and truncation of RGC repair following damage. Taken together, these data highlight a temporally conserved population of damage induced senescent immune cells that contribute to a complete and robust regenerative response in the zebrafish and provide new potential therapeutic targets to induce retina regeneration in mammals and humans.

## Acknowledgements

The authors would like to thank the Hitchcock Lab for their generous donation of the 4c4 antibody used in these experiments. They would also like to thank Qiang Guan and Jack Hollander for their assistance with the maintenance of the fish populations used in these experiments.

## Supplemental Figures

**Supplementary Figure 1.**
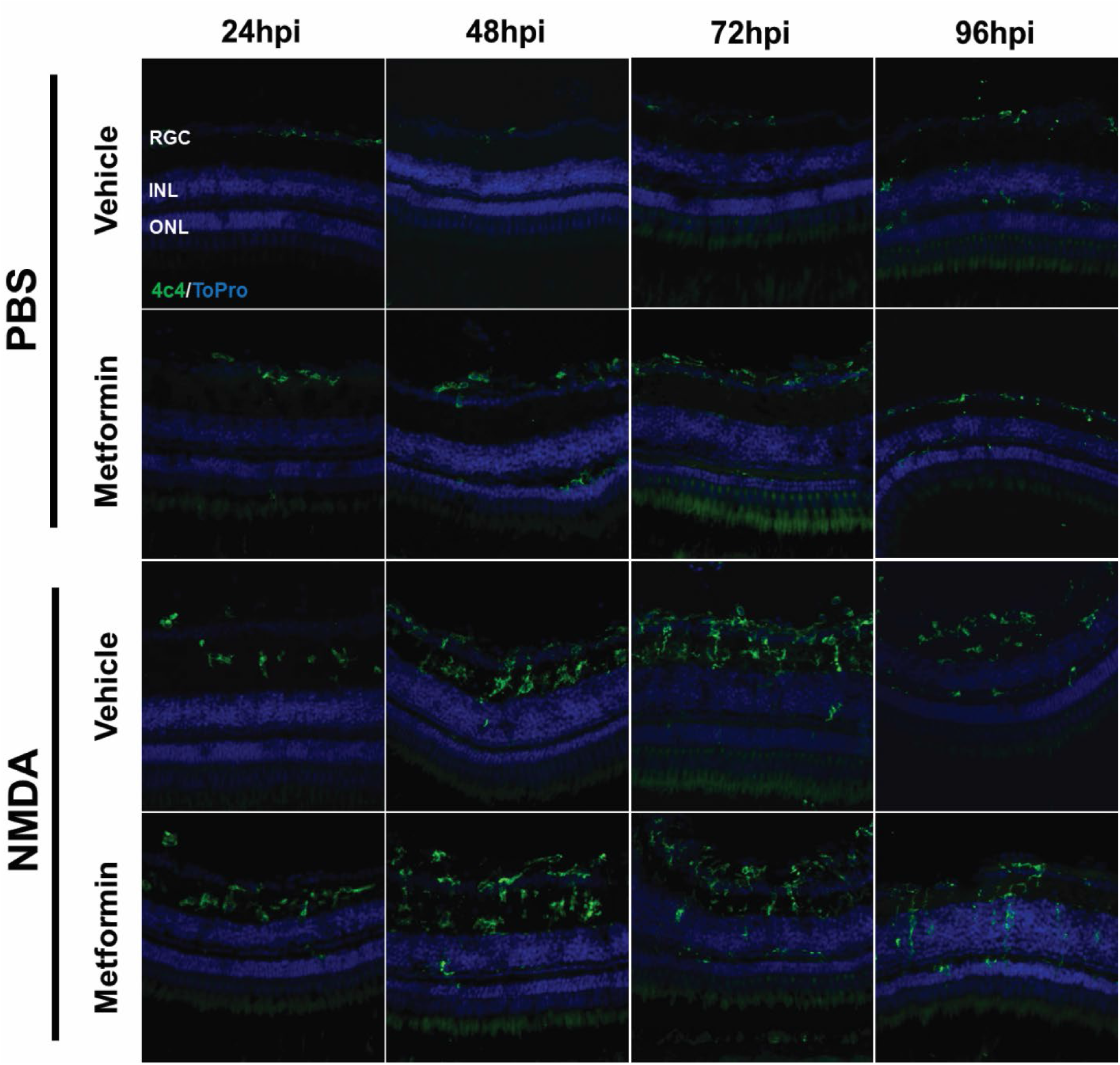
Metformin treatment does not impact microglial dynamics post NMDA damage. Wildtype AB zebrafish were intravitreally injected with 0.5uL of either 100mM NMDA or PBS, and then kept in tanks containing either 100uM of Metformin or standard tank water. Eyes were collected and sectioned and stained at early stages (24-96hpi) post damage to capture the inflammatory response post damage. Retinas that received Metformin treatment immediately following damage showed no significant difference in the number of 4c4^+^ macrophages/microglia (p>0.05, n=5).

**Supplementary Figure 2.**
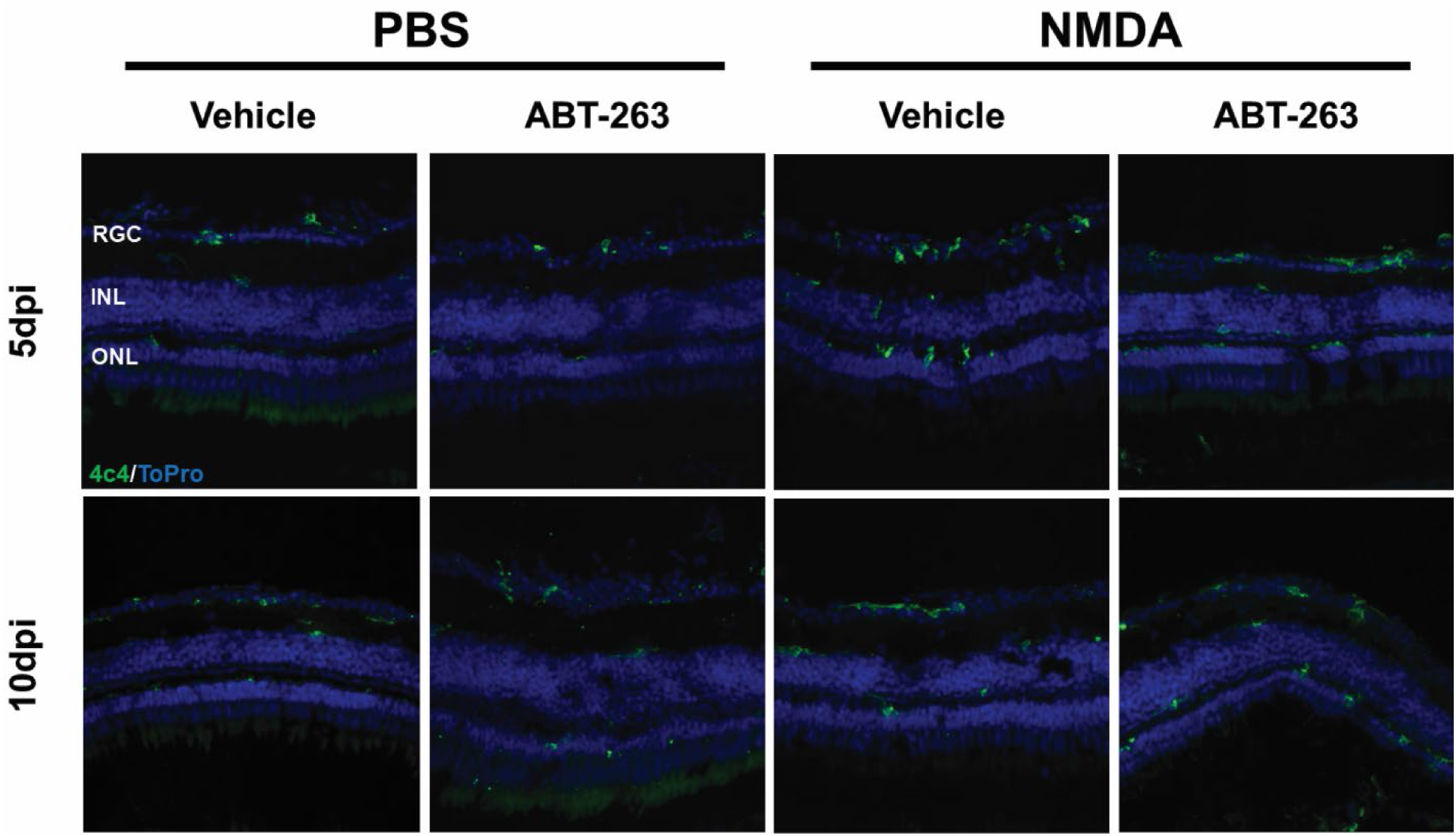
ABT-263 treatment does not impact microglial dynamics post NMDA damage. Wildtype (AB) zebrafish were intravitreally injected with 0.5uL of either 100mM NMDA or PBS, and subsequently intravitreally injected with 0.5uL of 30uM ABT-263 or vehicle control. Retinas receiving ABT-263 treatment showed no difference in the number of 4c4^+^ macrophages/microglia at 5 and 10dpi compared to untreated eyes (p>0.05, n=5).

**Supplementary Figure 3.**
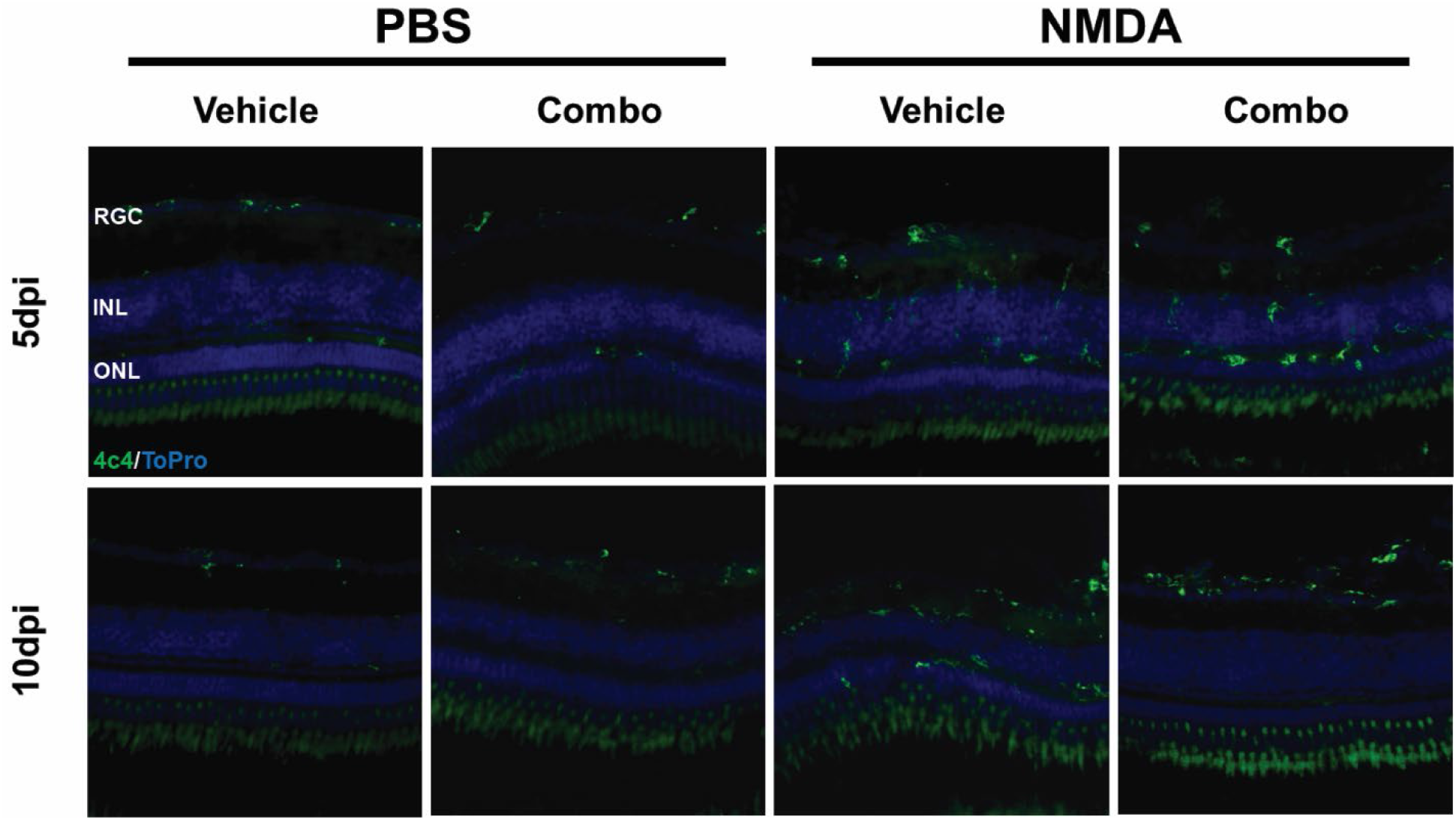
Combination senolytic treatment does not impact microglial dynamics post NMDA damage. Wildtype AB zebrafish were intravitreally injected with 0.5uL of either 100mM NMDA or PBS, and subsequently treated with a combination of Metformin and ABT-263 or vehicle control. Retinas that received the combination of senolytics showed no significant difference in the number of 4c4^+^macrophages/microglia at 5 and 10dpi compared to retinas that did not receive senolytic treatment (p>0.05, n=5).

**Supplementary Figure 4.**
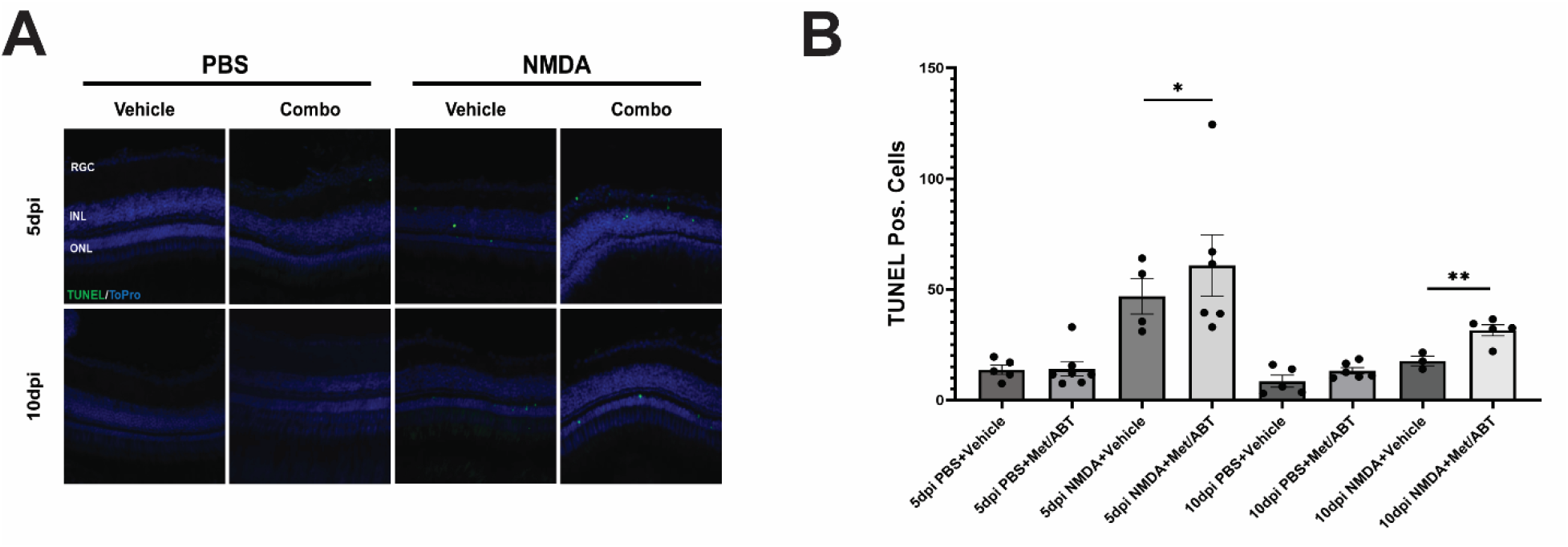
Senolytic treatment increases TUNEL^+^ cells after NMDA damage. Wildtype AB zebrafish were intravitreally injected with 0.5uL of either 100mM NMDA or PBS, and subsequently treated with a combination of Metformin and ABT-263 or a vehicle control. (A-B) Retinas that were treated with NMDA damage showed increased numbers of TUNEL^+^ cells across the whole retina, and combination senolytic treatment further increased the number of TUNEL^+^ cells in the retina at 5dpi (p<0.05, n=4-7) and 10dpi (p<0.005, n=3)

